# Indirect Deuterium Displacement Exchange Imaging for Non-invasive High-Resolution CSF Production Mapping

**DOI:** 10.1101/2025.10.20.683579

**Authors:** Debolina De, Shelei Pan, Helia Hosseini, Nicholas Bateman, Aliakbar Daemi, Joel R. Garbow, James D. Quirk, Jennifer M. Strahle, Arash Nazeri

## Abstract

Cerebrospinal fluid (CSF) production is central to brain homeostasis, yet existing measurement techniques are invasive, technically demanding, and confounded by intracranial perturbations. Here, we introduce indirect deuterium displacement exchange imaging (CSF-iDDxI), a noninvasive MRI approach for mapping CSF production *in vivo*. The method leverages intravenously infused deuterium oxide (D2O), which crosses the blood–CSF barrier, replaces existing H2O in CSF, producing concentration-dependent attenuation of proton (^1^H) signal within CSF spaces. Using high-resolution 3D balanced steady-state free precession MRI in rats, we demonstrate robust and spatially widespread D2O-induced CSF signal loss that is selectively suppressed by acetazolamide, a carbonic anhydrase inhibitor known to suppress CSF production by the choroid plexus. Dynamic imaging during intravenous D2O infusion further revealed that cortical parenchymal signal changes were unaffected by acetazolamide, confirming specificity to CSF production. Exploratory kinetic modeling estimated rapid CSF water turnover (k ≈ 0.09 min^-1^) under physiological conditions, reduced to k ≈ 0.031 min^-1^ with acetazolamide suppression, consistent with prior isotope tracer studies. Together, these findings establish CSF-iDDxI as a sensitive, pharmacologically validated tool for quantifying CSF production and turnover.

## Introduction

Cerebrospinal fluid (CSF) plays a central role in maintaining brain homeostasis by supporting waste clearance and intracranial pressure regulation^1^. CSF is primarily produced by the choroid plexus within the ventricular system, where a specialized epithelium forms the blood–CSF barrier (BCSFB)^2,3^, with possible extra-choroidal contributions from transcapillary and interstitial exchange^3^. Once secreted, CSF flows from the ventricles into the subarachnoid space, where it circulates around the brain and spinal cord. CSF penetrates the brain parenchyma via perivascular spaces, facilitating glymphatic clearance of metabolic by-products and proteins^1,2,4^. As the initiating driver of this cascade, CSF production serves as a key regulator of neurofluid homeostasis and waste clearance^1,5–7^. Disruption of this tightly regulated process has been implicated in hydrocephalus^8–11^, aging and age-associated neurodegeneration (e.g., Alzheimer’s disease)^12–15^, and idiopathic intracranial hypertension^16,17^.

Quantifying CSF production noninvasively thus represents a crucial step toward understanding how its dysregulation drives brain pathology and how targeted modulation might restore neurofluid homeostasis. Several methodologies exist for measuring CSF production, each with inherent advantages and limitations^18^. Direct measurements employing catheter-based quantification of CSF efflux from the lateral ventricles following surgical occlusion of the fourth ventricle yield precise estimates of ventricular choroidal CSF secretion^13^. Indirect methods typically involve infusing tracers or contrast agents into the ventricular system and measuring their dilution over time^18–20^. However, these approaches are susceptible to inaccuracies due to tracer leakage and incomplete mixing across CSF compartments^18^. Furthermore, these approaches are invasive, technically challenging, and can perturb intracranial pressure, potentially confounding the measurement of CSF production. As a non-invasive alternative, MRI-based approaches targeting CSF production and blood-to-CSF water exchange at the BCSFB have emerged as a promising in strategy^21^. Arterial spin labeling (ASL) MRI with ultra-long echo times has been recently introduced as a non-invasive indirect method to measure blood-to-CSF water exchange across the BCSFB, leveraging the long T2 of CSF to isolate signal from labeled water entering the ventricular space^21–23^. However, this approach is inherently complex, integrating multiple physiological parameters, relying on modeling assumptions, limited by low spatial resolution, and insensitive to slow or cumulative changes in CSF production dynamics.

To address the limitations of existing methods, we developed indirect <u>d</u>euterium <u>d</u>isplacement e<u>x</u>change imaging for mapping CSF production (CSF-iDDxI) using intravenously infused deuterium oxide (D2O, heavy water; **Fig. 1**). This approach provides the imaging analog of the water-exchange paradigm originally pioneered by Bering in the 1950s^24^. As a stable isotopic analog of H2O, D2O is functionally equivalent to water in its transport properties and distribution across physiological compartments through aquaporins^25,26^, making it an ideal tracer for studying water dynamics *in vivo*.

**Figure 1.**
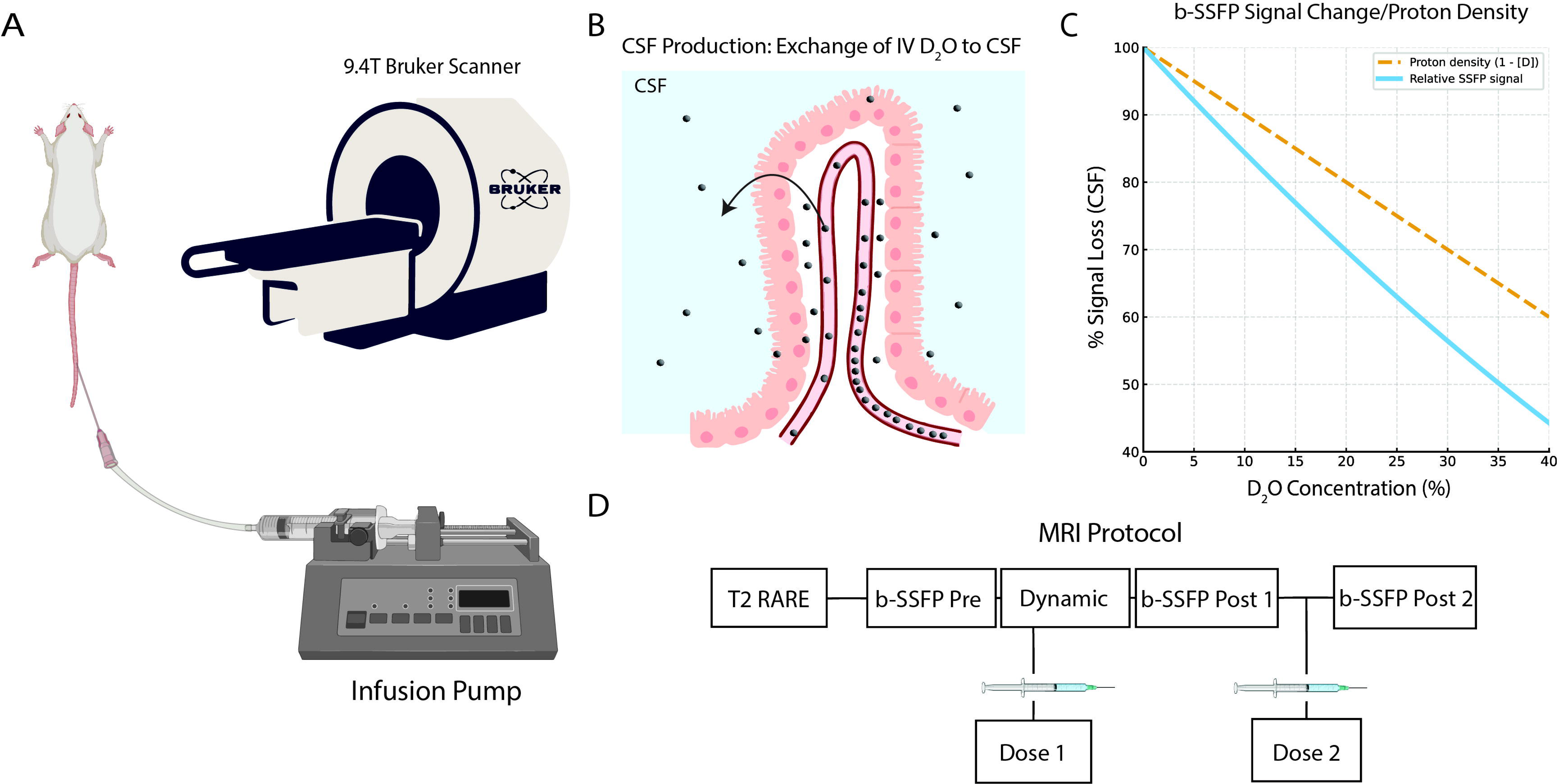
Experimental framework for indirect deuterium displacement exchange imaging for mapping CSF production (CSF-iDDxI) in rats. (A) Setup for intravenous (IV) infusion of deuterium oxide (D2O) during 9.4T small-animal MRI. D2O was delivered via tail vein catheter using a programmable infusion pump. (B) Schematic of D2O exchange into cerebrospinal fluid (CSF) across the blood-CSF-barrier (BCSFB). Following intravenous infusion, D2O traverses the choroid plexus epithelium and enters the CSF spaces as newly secreted CSF. (C) Relationship between D2O concentration and CSF signal change in b-SSFP. Proton density decreases linearly with increasing D2O fraction, while the b-SSFP signal exhibits a steeper decline reflecting combined effects of proton dilution and relaxation changes. (D) Imaging and infusion protocol. After anatomical T2-weighted scans (2D T2-rapid acquisition with relaxation enhancement [RARE]) and baseline balanced steady-state free precession (b-SSFP), the first intravenous D2O infusion was administered during dynamic deuterium displacement imaging (‘Dynamic’). Subsequent b-SSFP acquisitions were used to quantify CSF signal attenuation after the first and second D2O infusions, respectively.

D2O can be indirectly detected on conventional proton (^1^H) MRI through a concentration-dependent signal loss caused by displacement of native tissue water^27,28^. This principle has been previously demonstrated for assessing cerebral perfusion, where intravascular D2O infusion led to measurable ^1^H signal attenuation in the brain parenchyma^27,28^. In contrast to brain parenchyma, where signal changes arise from both intravascular D2O and D2O that has crossed the blood–brain barrier (BBB), CSF lacks a blood compartment; thus, any D2O-related signal loss in CSF reflects water molecules that have exchanged into the compartment.

Building on this foundation, we utilized D2O-induced signal loss for CSF labeling: following infusion, D2O circulates systemically and crosses the BCSFB, resulting in newly secreted CSF with a D2O concentration that closely reflects that of plasma. This replacement of H2O by D2O enables visualization of CSF production as a progressive reduction in ^1^H signal intensity within the ventricular and extra-ventricular CSF spaces, proportional to the influx of D2O. Given the relatively small size of CSF spaces in murine models, evaluation of D2O-induced signal attenuation with CSF-iDDxI requires imaging techniques with high spatial resolution, strong signal-to-noise ratio (SNR), and insensitivity to flow-related artifacts. To address these constraints, we employ high-resolution three-dimensional balanced steady-state free precession (3D b-SSFP), which provides sharp fluid–tissue contrast and is well suited for detecting subtle changes in CSF composition without partial volume effects. We applied this approach in a rat model and demonstrated that D2O infusion produced robust signal loss within the CSF. To validate the approach, we used acetazolamide (ACZ), a carbonic anhydrase inhibitor that suppresses CSF production. ACZ markedly attenuated D2O-induced CSF signal loss, while cortical dynamics were unaffected, indicating that the effect reflects specific suppression of CSF production rather than nonspecific consequences of diuresis or perfusion. Exploratory kinetic modeling further confirmed that CSF water turnover is rapid under physiological conditions and slows markedly with acetazolamide. Together, these results establish a noninvasive framework for imaging CSF production and its pharmacological modulation *in vivo*.

## Results

### D2O-induced global CSF signal loss is attenuated by acetazolamide

Using the CSF-iDDxI paradigm (**Fig. 1**), voxel-wise maps of CSF secretion were derived by comparing pre- and post-infusion b-SSFP signal intensities within CSF spaces. D2O was infused twice to confirm reproducibility of the CSF-iDDxI measurements across repeated trials. Rats were assigned to three groups: D2O infusion, H2O infusion as control, and D2O infusion following acetazolamide pretreatment (D2O/ACZ) to suppress CSF production. Group-averaged signal loss maps showed widespread attenuation across the CSF spaces with D2O, minimal change with H2O, and intermediate values with D2O/ACZ (**Fig. 2A**). To quantify these effects, global CSF signal changes were calculated across the entire CSF compartment. Global CSF signal decrease (averaged across the CSF compartment) was larger in the D2O group (infusion 1: 9.3% ± 3.1%; infusion 2: 6.4% ± 2.3%) compared to the H2O group (infusion 1: 1.2% ± 2.3%, p = 0.0023; infusion 2: 0.7% ± 2.4%, p = 0.0023; **Fig. 2B**). The D2O/ACZ group showed intermediate signal loss (infusion 1: 5.5% ± 1.6%; infusion 2: 3.7% ± 1.3%), significantly lower than D2O (infusion 1: p = 0.042; infusion 2: p = 0.019) but higher than H2O (infusion 1: p = 0.0007; infusion 2: p = 0.008; **Fig. 2B**). Thus, a single intraperitoneal dose of ACZ (40mg/kg) suppressed D2O-induced signal loss by 46% following the first infusion and 48% following the second. Relative to the first infusion, the second infusion produced 31% less signal loss in the D2O group and 33% less in the D2O/ACZ group. This likely reflects that the first infusion introduces D2O into a naïve CSF water pool with minimal clearance or back exchange across the BCSFB, whereas by the second infusion, the CSF is already partially deuterated, leading to enhanced D2O efflux and exchange across the barrier.

**Figure 2.**
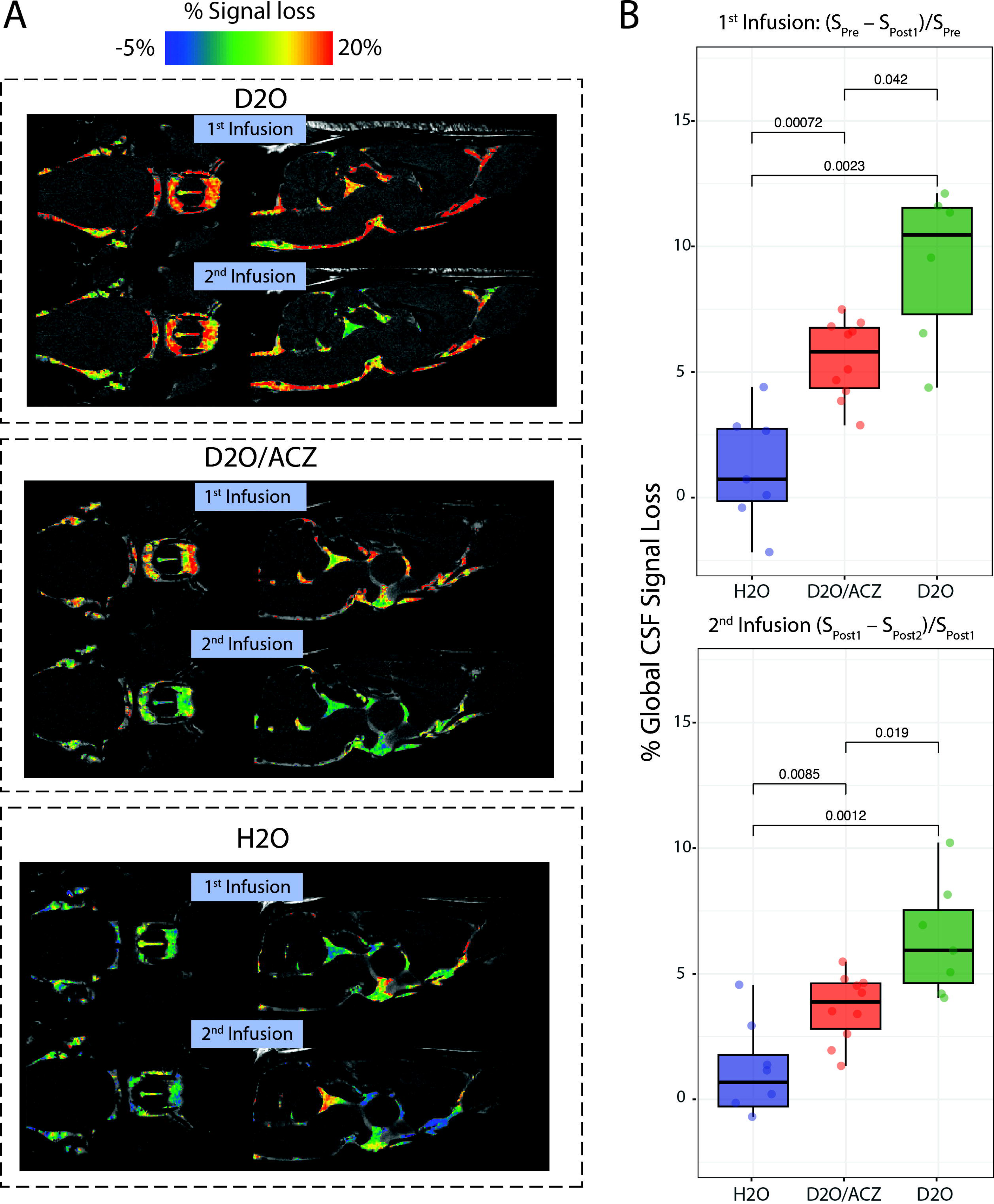
D2O-induced global CSF signal loss and pharmacological suppression by acetazolamide. (A) Representative color-coded maps of % CSF signal loss in rats infused with D2O (top), D2O following acetazolamide pretreatment (middle), or H2O vehicle (bottom). Percent signal change is overlaid on anatomical images for the first and second infusion periods. (B) Quantification of global CSF signal loss across groups. Top: signal loss following the first infusion. Bottom: signal loss following the second infusion. *p-values* indicate results of between group non-parametric statistical comparisons (Wilcoxon rank-sum test).

### Region-of-interest analysis reveals widespread attenuation of D2O-induced CSF signal loss by acetazolamide

To further investigate the spatial distribution of CSF production, a region-of-interest (ROI) analysis was performed across five ventricular and eight cisternal CSF compartments (**Fig. 3A**). Percent signal changes were calculated using the median signal intensity in each ROI before and after each infusion. Similar to the global CSF signal changes, Jonckheere–Terpstra tests revealed a significant trend (D2O > D2O/ACZ > H2O) across nearly all ROIs for both infusions (**Fig. 3B**). Compared with the H2O group, the D2O group showed greater signal loss across most ROIs (21/26, 81%, one-tailed p < 0.05). In contrast, pairwise comparisons involving the D2O/ACZ group revealed fewer significant differences (one-tailed p < 0.05) relative to either H2O (11/26, 42%) or D2O (7/26, 27%). These nonsignificant effects lacked a clear ventricular–cisternal distribution. They likely reflect smaller effect sizes within individual ROIs and reduced between group difference-to-noise compared with global CSF signal changes. Together, the results indicate that, by the time of post-infusion imaging, ACZ exerts a diffuse, rather than regionally selective, attenuation of D2O-induced CSF signal loss.

**Figure 3.**
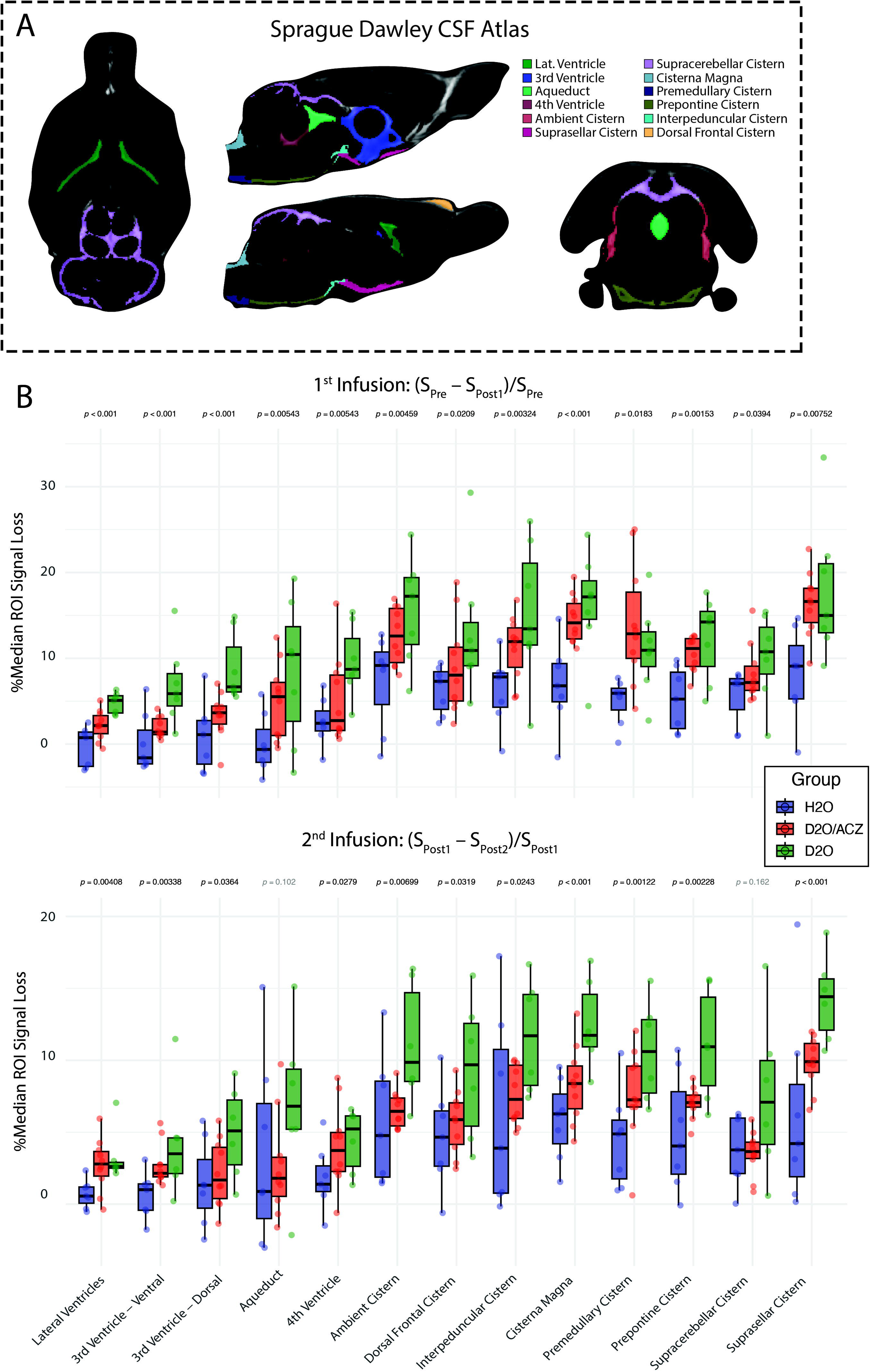
Regional CSF signal loss following D2O infusions. (A) Atlas-based segmentation of major ventricular and cisternal CSF compartments in the adult Sprague-Dawley rat brain. B) Quantification of median CSF signal loss across atlas-defined regions of interest (ROIs) following intravenous infusions. Statistical comparisons were performed using the Jonckheere–Terpstra test with the hypothesized order D2O > D2O/ACZ > H2O (one-tailed).

### Acetazolamide does not alter D2O-induced signal loss in the cerebral cortex

Indirect dynamic deuterium displacement imaging was applied to assess D2O-induced signal attenuation in the cerebral cortex, calculated as percent signal loss before and after infusion. As expected, D2O produced a robust negative contrast during infusion. In both the D2O and D2O/ACZ groups, cortical signal reached a plateau after infusion cessation (frame >106, ∼14 minutes). We hypothesized that if the CSF effects were driven by nonspecific consequences of diuresis or altered perfusion rather than by choroid plexus–mediated CSF production, cortical D2O-induced signal loss would also be attenuated by ACZ. Cortical signal loss was significantly greater in both the D2O and D2O/ACZ groups compared to the H2O group, with no significant difference between the D2O and D2O/ACZ groups (**Fig. 4**). These results confirm that the attenuation of CSF signal loss by ACZ reflects specific suppression of CSF production, rather than global alterations in brain water dynamics or nonspecific diuretic effects.

**Figure 4.**
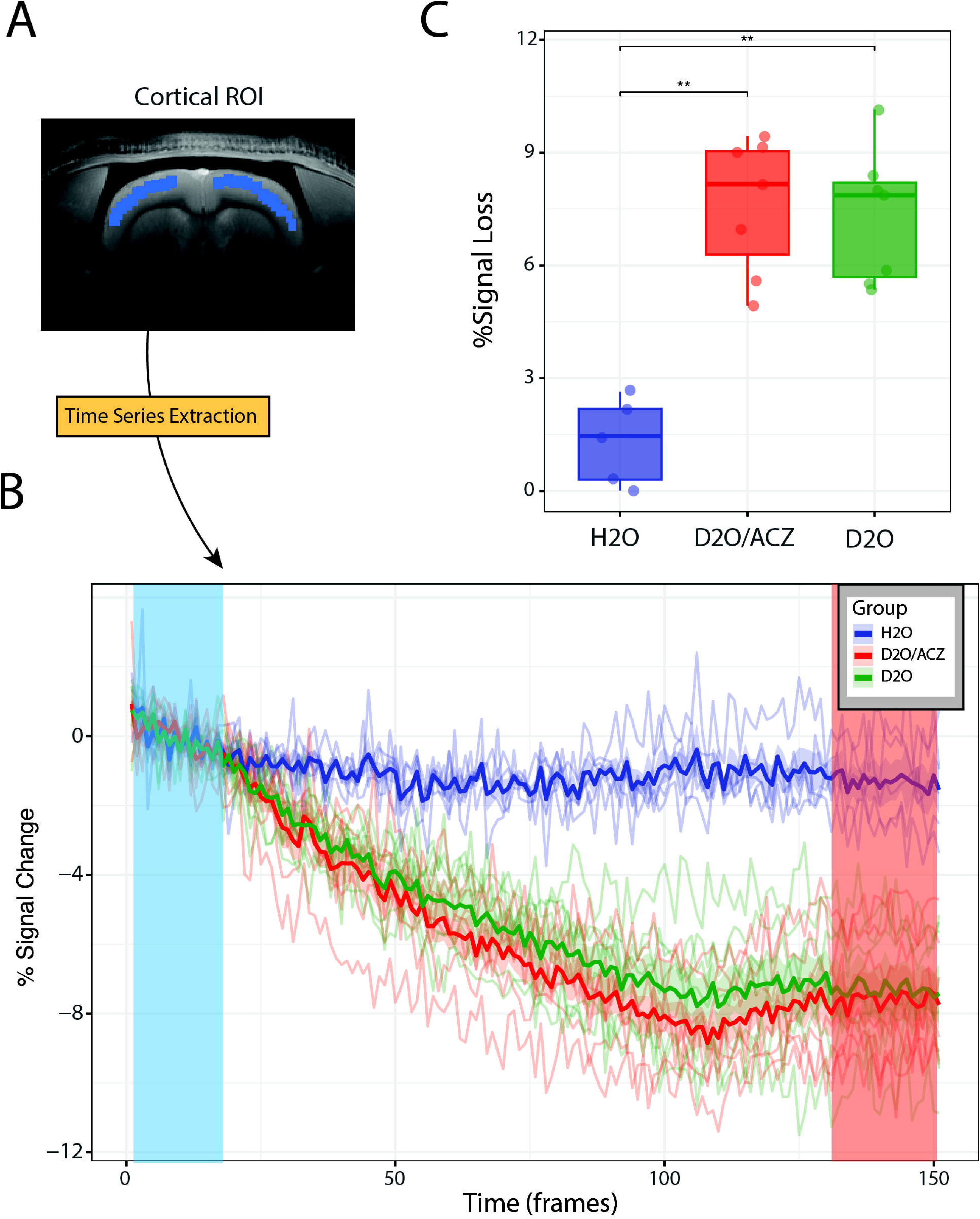
Cortical dynamics of deuterium displacement during infusion. (A) Cortical region-of-interest (ROI) mask used for time-series extraction from dynamic deuterium displacement imaging. (B) Cortical signal trajectories before, during, and after intravenous infusion. Baseline (blue) was defined as the 2nd–15th volumes prior to D2O/H2O infusion, and the post-infusion window (orange) as the final 20 volumes. Cortical signal declined progressively with D2O infusion, independent of acetazolamide pretreatment. (C) Quantification of relative cortical signal loss, expressed as percent change between baseline and post-infusion periods. Both D2O and D2O/ACZ groups exhibited significant attenuation compared to H2O controls, with no difference between D2O and D2O/ACZ (**p < 0.01).

### Exploratory kinetic modeling demonstrates fast CSF water turnover rate

To quantify CSF D2O kinetics, we first characterized the plasma input D2O concentration (C_p_) from the superior sagittal sinus using a three-segment piecewise model with fixed breakpoints defined by the first D2O infusion paradigm (baseline before infusion, downslope during infusion, plateau after infusion; sampling interval = 7.95 s; n = 4; Fig. 5). Dynamic deuterium displacement imaging signal changes were converted to plasma D2O concentration (C_p_) using the gradient-echo signal model, which incorporates both proton density replacement and T1 relaxation effects (Methods; Equation 3). The group-averaged C_p_ trajectory exhibited a linear downslope of −0.56 %/min and reached a ∼6.7% attenuation at plateau. The piecewise model closely captured the group mean (R^2^ > 0.95). To translate these empirical dynamics into physiological turnover constants, we applied the convolution model (Methods; Equations 4 and 5; Fig. 5) using D2O concentrations derived from the b-SSFP signal equation. Specifically, the mean percent signal losses relative to the H2O vehicle after the first infusion (D2O – H2O: 8.1%; D2O/ACZ – H2O: 4.3%) were converted to CSF D2O concentrations (C_CSF_(T)) of 5.09% and 2.68%, respectively. Using the piecewise-linear plasma input function and measured CSF endpoint concentration, the nonlinear solver yielded a CSF water turnover constant of k ≈ 0.09 min^-1^ in the D2O group and k ≈ 0.031 min^-1^ in the D2O/ACZ group

**Figure 5.**
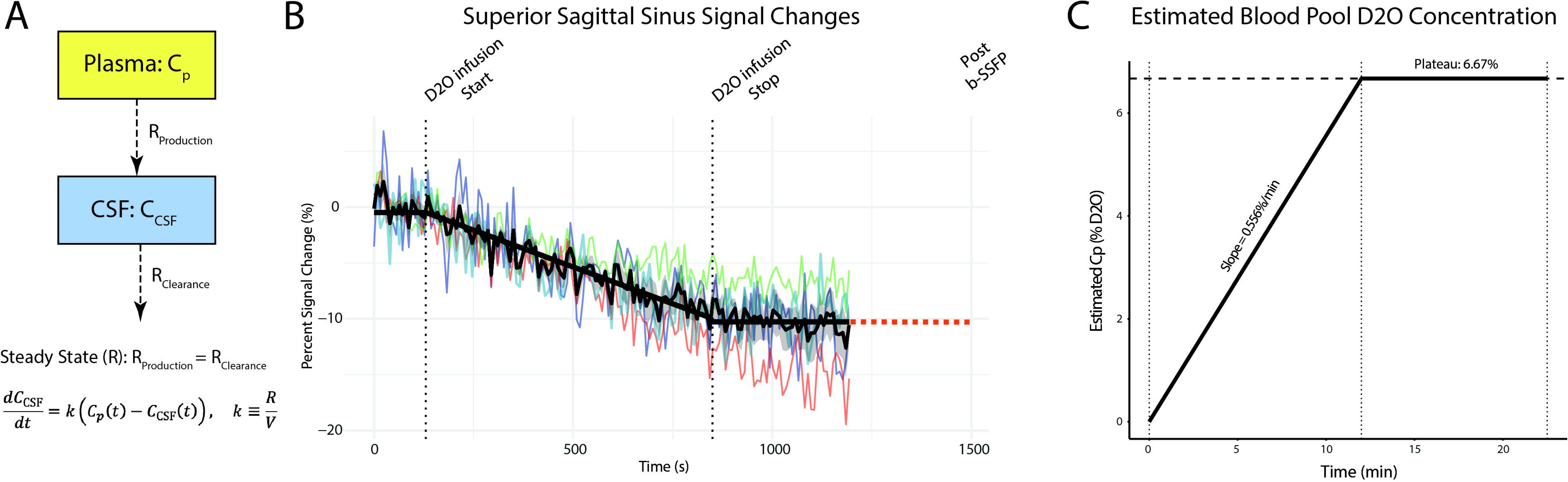
Kinetic modeling framework for CSF production analysis. (A) One-compartment exchange model describing the relationship between plasma (C_p_) and CSF (CCSF) D2O concentrations. At steady state, CSF production equals clearance (R_Production_ = R_Clearance_), as CSF volume remains unchanged. CSF turnover is defined by the rate constant k=R/V, where R is the production/clearance rate and V is CSF volume. (B) Time-resolved signal changes in the superior sagittal sinus, used as a proxy for D2O-related signal change in the blood pool. Thin colored traces represent individual animals; the bold black line shows the group mean ± standard error. Dotted vertical lines denote the start and stop of intravenous D2O infusion, followed by post-infusion b-SSFP imaging. The dashed orange line indicates the presumed steady-state plasma D2O concentration at the time of CSF imaging (T). (C) Estimated plasma D2O concentration over time, modeled from venous signal changes. The concentration increases linearly during infusion (slope = 0.556 %/min) until reaching a plateau of 6.67%, which is maintained during the post-infusion steady state.

## Discussion

Our findings establish intravenously infused D2O combined with high-resolution b-SSFP MRI as a noninvasive strategy for probing CSF production dynamics *in vivo*. We demonstrate that systemic D2O administration produces robust and spatially widespread attenuation of CSF signal, consistent with incorporation of D2O into newly secreted fluid across the BCSFB. Critically, this effect was markedly suppressed by ACZ, a carbonic anhydrase inhibitor that reduces choroid plexus secretion. To capture temporal dynamics, we complemented high-resolution CSF-iDDxI with dynamic deuterium displacement imaging during, tracking D2O-induced signal loss in both brain parenchyma and the superior sagittal sinus (blood pool) during and after the first infusion. In contrast, cortical D2O-induced signal changes were unaffected by ACZ, confirming that the observed CSF effects reflect targeted suppression of secretion rather than nonspecific alterations in perfusion or water distribution.

Heavy water (D2O) can be detected *in vivo* with MRI either directly with a dedicated deuterium (^2^H) coil or indirectly with a conventional ^1^H coil. In the seminal studies direct ^2^H imaging was first applied in early studies of perfusion^29^ and has more recently been extended to metabolic imaging^30^, but its relatively low SNR constrains achievable spatial resolution^31^. For indirect deuterium imaging, given that D2O is invisible to ^1^H coil, its displacement of H2O leads to a reduction in ^1^H signal intensity, providing an indirect readout of tracer dynamics^27,28,32^. Indirect deuterium imaging has also been used for imaging cerebral perfusion, by characterizing D2O-induced signal loss on 1H coil^27,28^.

Whereas earlier studies employed dynamic, low-resolution 2D acquisitions to track perfusion-related signal changes, we employed 3D b-SSFP at 110 µm^3^ resolution for CSF-iDDxI to enable high-resolution mapping of CSF signal changes. Such resolution is essential in murine models given the small intracranial CSF spaces and the need to minimize partial volume effects. In addition, signal changes in CSF-iDDxI represent exchanged D2O, as vascular partial volume contribution is negligible within high intensity CSF. This differs from prior indirect deuterium imaging studies, where signal changes reflect a combination of intravascular D2O and D2O that had crossed the BBB into interstitial and intracellular compartments.

3D b-SSFP is particularly well-suited for this application, combining rapid volumetric acquisition and high SNR efficiency with relative insensitivity to flow-related artifacts that typically confound CSF imaging. However, the long acquisition time (4–5 minutes per phase advance, with at least two phases) precludes fully dynamic imaging at this resolution, and the low parenchymal SNR limits detection of tissue signal changes.

However, for translation to larger animals and humans, where CSF spaces are larger, whole-brain acquisitions at submillimeter resolutions can be achieved with temporal resolutions <1 minute using advanced acceleration techniques^33^.

We found that pretreatment with a single dose of ACZ (40 mg/kg) suppressed D2O-induced signal loss in CSF production by ∼47%. The suppressive effect of ACZ on CSF secretion is well established, thereby serving as a pharmacological validation for CSF-iDDxI. Previous studies using alternative approaches for measuring CSF production (e.g., ventriculo-cisternal perfusion) have reported comparable ACZ-induced reductions (40–57%) across various animal models^34–37^. ACZ-induced changes were confined to CSF signal, as cortical parenchymal ROIs showed no difference between the D2O and D2O/ACZ groups on dynamic deuterium displacement imaging. These findings indicate that ACZ’s effect is specific to CSF production and not attributable to nonspecific diuretic effects or perfusion-related changes. Overall, CSF-iDDxI establishes a platform for quantitative measurement of CSF production *in vivo* and for evaluating drugs that suppress or enhance secretion, as well as diseases with suspected disturbances in CSF production.

Follow-up ROI analysis revealed no spatially distinct effects of ACZ. This presumably reflects equilibration of water composition across CSF spaces within 10–15 minutes of infusion. This is likely facilitated by mixing effects from cardiorespiratory-driven CSF flow, which enhance effective CSF diffusivity^38,39^. Resolving potential choroidal versus extrachoroidal sources of CSF production, or detecting spatial gradients, may require higher temporal resolution or CSF-iDDxI performed during continuous intravenous D2O infusion. Similarly, intra-arterial D2O administration could reveal lateralized as well as choroidal versus extra-choroidal spatial gradients. CSF-iDDxI–based spatial mapping of CSF production may have translational relevance, particularly in obstructive hydrocephalus, where ventricular entrapment prevents newly secreted CSF from circulating beyond the site of obstruction.

Our exploratory estimates of CSF kinetics demonstrated CSF water turnover constant of k ≈ 0.09 min^-1^ under physiological conditions and k ≈ 0.031 min^-1^ with ACZ-mediated CSF production suppression. These values align with prior evidence from tritiated water (HTO) studies in animals, which have shown rapid equilibration between plasma and CSF within 5–10 minutes after bolus injection^40,41^, reflecting fast CSF turnover dynamics comparable to those observed here. Of note, CSF water turnover rates derived from our approach are not directly comparable to production rates reported with earlier methods. For instance, Liu et al. reported a CSF production rate of 133.2 nL/min under isoflurane anesthesia (assuming V_CSF_ ≈ 36 µL in mice; k = 0.0037 min^-1^)^13^. This estimate, however, does not account for contributions from the fourth ventricle or extra-choroidal sources, and more fundamentally represents net CSF efflux through a lateral ventricular cannula rather than steady-state CSF water turnover rate.

Several limitations warrant consideration. First, the relatively long acquisition times of high-resolution 3D b-SSFP preclude fully dynamic mapping of CSF production, limiting the ability to resolve rapid fluctuations or compartment-specific gradients. Future large-animal studies leveraging advanced acceleration techniques may enable dynamic, high-resolution CSF-iDDxI with improved temporal resolution and translational relevance.

Second, in small-animal preclinical MRI, surface coil–related receive bias can heighten the impact of minor head motion, leading to compromised signal uniformity and subtle motion-related signal fluctuations. Securing the head with extra caution is therefore essential in small animals, while the use of multi-channel coils in clinical scanners should mitigate these effects. In this study, we utilized H2O as the vehicle for accounting for such signal drifts in our experiments. Third, we employed relatively large-volume isotonic saline infusions, a standard method in rodent fluid expansion studies^42,43^.

Although generally safe, such supraphysiologic loading can transiently alter hemodynamic and electrolyte balance and should be taken into account when interpreting the findings. Finally, our kinetic analysis should be viewed as proof of principle, as intravascular and CSF tracer concentrations were not measured simultaneously, and superior sagittal sinus values were used as a proxy for the blood pool. Given that arterial D2O concentrations during active D2O infusion are likely higher than those in the venous outflow, this approximation could lead to a slight overestimation of k. Moreover, high-resolution b-SSFP imaging required 4:20 per phase advance (8:40 for two), such that the CSF concentration time point (*T*) was effectively assigned to the end of the first acquisition. Future implementation of accelerated imaging strategies and translation to larger animal models will enable higher temporal resolution dynamic imaging, mitigating these limitations and permitting more accurate kinetic modeling.

Together, these findings establish CSF-iDDxI as a noninvasive imaging framework for quantifying CSF production and its pharmacological modulation *in vivo*. This approach may also be applied to disease models to examine pathological alterations in CSF production and turnover. For accurate kinetic modeling, future studies will need to incorporate imaging strategies that capture CSF and intravascular D2O signal dynamics simultaneously. Large-animal studies will be critical for testing the feasibility of the approach with lower intravenous D2O doses suitable for human translation. Ultimately, this framework provides a foundation for systematic evaluation of therapeutic strategies that alter CSF secretion and neurofluid dynamics.

## Methods

### Animals

All experiments were approved by the Institutional Animal Care and Use Committee at Washington University in St. Louis (Protocol ID: 24-0027). In total, 25 male Sprague– Dawley rats (Charles River Laboratories, Wilmington, MA), weighing between 550 and 650 g and aged 4–6 months, were used in this study.

### Experimental Design

Rats were randomly assigned to one of three groups: (i) D2O group (n = 7), which received intravenous deuterium oxide; (ii) negative control H2O group (n = 8), which received intravenous H2O as the vehicle; and (iii) D2O-ACZ group (n=10), which was pretreated with an intraperitoneal injection of ACZ (40mg/kg) immediately prior to MRI to suppress CSF production^44^. All animals received two intravenous infusions of normal saline prepared in either D2O or H2O. Each infusion was warmed to 35–37 °C, administered at 15 mL/kg, and delivered over 12 minutes via tail vein using an infusion pump. All animals receiving D2O remained viable for the entire study period until planned euthanasia, with no premature mortality (4–6 months of follow-up).

### MRI Acquisition

All MRI experiments were performed at the Small Animal Magnetic Resonance Facility of the Mallinckrodt Institute of Radiology at Washington University in St. Louis. Anesthesia was induced using 3-4% isoflurane. Once inside the scanner the isoflurane was reduced to 1.8-2.5% and the animal was fixed using a bite bar to the MR bed. The breathing rates were monitored using a respiration pillow sensor (SA Instruments Inc., Stony Brook, USA) and was in the range of 45-60 bpm. The MR bed kept at 50°C. Each scanning session lasted for a total of 1.25 hours. The first infusion commenced 25–30 minutes after scanning began. The second infusion started ∼17 minutes after the first infusion.

All images were acquired using a 9.4 T Bruker Biospin preclinical MRI system (Bruker, Billerica, MA) equipped with a dedicated 4-channel rat head coil. An axial 2D T2-weighted rapid acquisition with relaxation enhancement (T2-RARE) sequence was used to acquire anatomical reference images to perform anatomical brain segmentation. The T2-RARE sequence parameters were field of view (FOV): 100 x 100 x 300 µm^3^; RARE factor: 8; echo time (TE): 39 ms; repetition time (TR): 7200 ms. A high-resolution 3D b-SSFP sequence was utilized to capture deuterium displacement – related signal loss in the CSF. The b-SSFP parameters: flip angle: 50°; TE: 2.35 ms; TR: 4.69 ms; voxel-size: 110 x 110 x 110 µm^3^; acquisition time: 4:21 (per phase advance). This sequence was repeated with phase advances of 0° and 180°. They were performed before and after each infusion. To evaluate signal-loss dynamics in brain parenchyma (cerebral cortex) and neurovasculature (superior sagittal sinus), a subset of animals (n = 19; D2O: 7; H2O: 5; D2O/ACZ: 7) underwent indirect dynamic deuterium displacement imaging using a multi-slice 2D gradient echo sequence: number of volumes: 151; temporal resolution: 7.95s; Flip angle: 15°; TE: 3.6; TR: 53 ms; in-plane resolution: 93 x 80 µm^2^; slice thickness: 2 mm; number of slices: 4. D2O infusion began approximately 2 minutes after the start of dynamic deuterium displacement image acquisition (volume ∼17-18) and dynamic imaging continued for ∼6 minutes after infusion was stopped. A large slice thickness was chosen to enhance SNR while maintaining high in-plane resolution and to minimize flow-related signal variations.

### Whole-brain Analysis of D2O-Induced CSF Signal Changes

High-resolution b-SSFP data were analyzed to evaluate D2O displacement in the CSF, quantified as percentage signal drop as a surrogate marker of CSF production. Banding artifacts in the b-SSFP images were corrected by merging phase-sensitive acquisitions (0° and 180°) and computing the square root of the sum of squares (SOS)^45^. Post-infusion images were co-registered to pre-infusion baselines using ANTs^46^. A manually segmented brain mask from T2-weighted images was used for skull stripping, followed by CSF segmentation using Atropos^47^. Percentage signal change maps were computed for two intervals: from pre-infusion to the first post-infusion timepoint, and from the first to the second post-infusion timepoint. To estimate global CSF displacement, mean percentage signal loss across the entire CSF compartment was computed at each timepoint using *fslstats* (part of FSL v6.0)^48^.

### Region-of-interest analysis of D2O-induced CSF signal changes

The brain mask was applied to the mean 3D b-SSFP image to obtain a skull-stripped brain volume. The b-SSFP images from all animals were used to create a population template via iterative nonlinear registration in ANTs (*antsMultivariateTemplateConstruction2.sh* script)^46^. A rat CSF atlas comprising thirteen ROIs was manually labeled in template space using ITK-SNAP (v4.2.0)^49^. The ROIs encompassed ventricular and cisternal compartments, including the lateral ventricles, ventral and dorsal third ventricle, cerebral aqueduct, fourth ventricle, ambient cistern, dorsal frontal cistern, interpeduncular cistern, cisterna magna, premedullary cistern, prepontine cistern, supracerebellar cistern, and suprasellar cistern. The atlas was then nonlinearly registered to each animal’s native space using the inverse warp fields generated during template construction. The median signal intensity was extracted from each ROI in native space, and percent signal changes were calculated for the pre– post1 and post1–post2 b-SSFP volumes.

### Indirect dynamic deuterium displacement imaging analysis

An ROI was manually delineated over the cerebral cortex using ITK-SNAP^49^. Mean signal intensity was extracted from the ROI for each volume using *fslstats*. To ensure image stability, the first two volumes were excluded. Pre-infusion signal was defined as volumes 3–22, and post-infusion signal as the final 20 volumes of the acquisition. The percentage signal loss was then calculated as the relative decrease from pre- to post-infusion values.

### Estimation of D2O concentration

D2O-related b-SSFP signal changes in CSF reflect both proton density replacement and relaxation effects (equation 1):

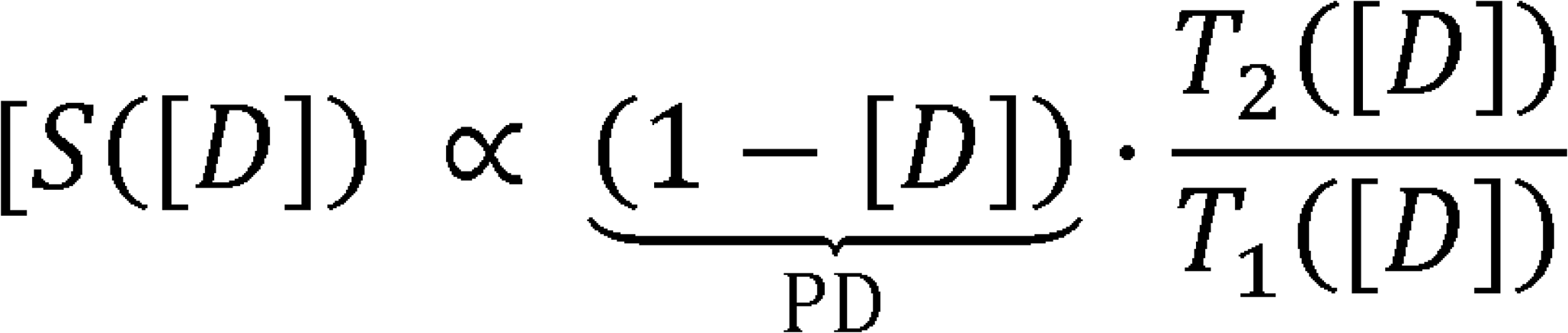

Relaxation rates were modeled as (equation 2):

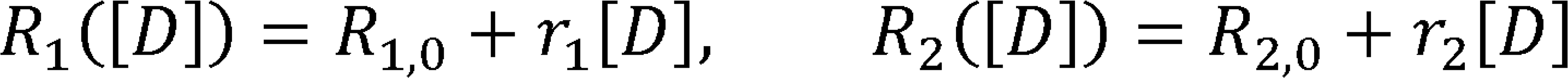

Baseline values were T1,0 = 3.3 s and T2,0 = 0.67 s for CSF at 9.4 T^50,51^, with relaxivities r1 = −0.23/s and r2 = -0.21/s per D2O fraction, obtained from aqueous phantom calibrations at 11.7T^28^. D2O-related blood signal changes in dynamic deuterium displacement imaging reflect a combination of reduced proton density and prolonged T1 relaxation, while T2 effects could be ignored due to the very short TE (equation 3):

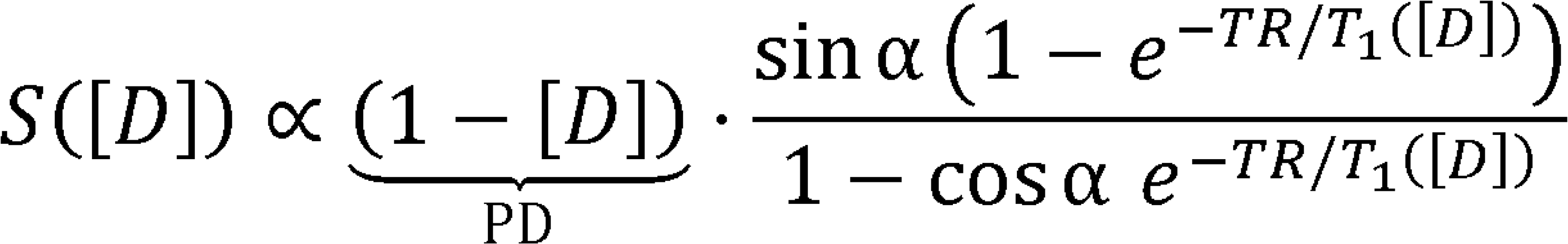

Here, (1−[D]) accounts for proton replacement, and T1([D]) reflects the T1 relaxation as of D2O. We used T1,0 = 2.44 s for blood at 9.4 T^52^ and r1 = −0.230/s^28^. Based on these equations, D2O concentrations in CSF (C_CSF_) and blood (plasma; C_p_) were estimated from DDxI b-SSFP images and dynamic deuterium displacement gradient echo images, respectively.

### Kinetic modeling for quantification of CSF production

We modeled CSF D2O kinetics under the assumption of steady state, such that CSF production and clearance are equal and total CSF volume remains constant during the experiment. Specifically, we hypothesized that: (i) signal changes in the CSF arise from the appearance of newly secreted CSF containing D2O; (ii) the D2O concentration of the newly formed CSF equals the instantaneous plasma D2O concentration (C_P_); and (iii) D2O in the blood is equilibrated between plasma and red blood cells, such that plasma concentration reflects the whole blood pool. At steady state, the time course of CSF D2O concentration (C_csf_), follows a one-compartment exchange model (equation 4):

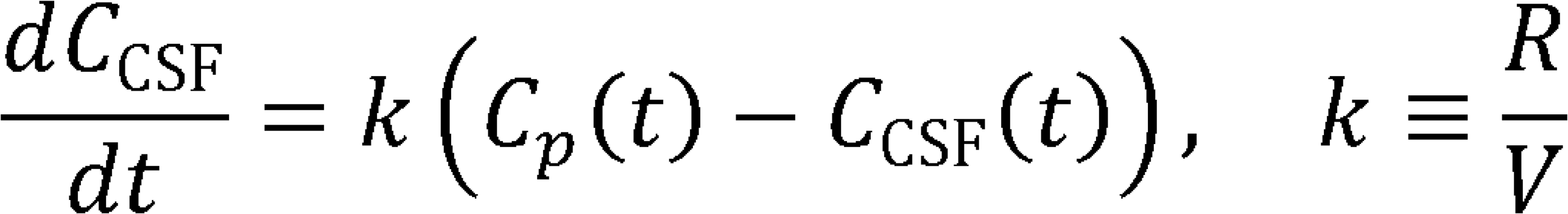

where k is turnover rate (1/min), representing the fraction of CSF volume replaced per unit time. Therefore, *k* is defined as CSF production rate (R; µL/min) divided by CSF volume (V, µL). The solution at time T (post D2O infusion b-SSFP) is (equation 5):

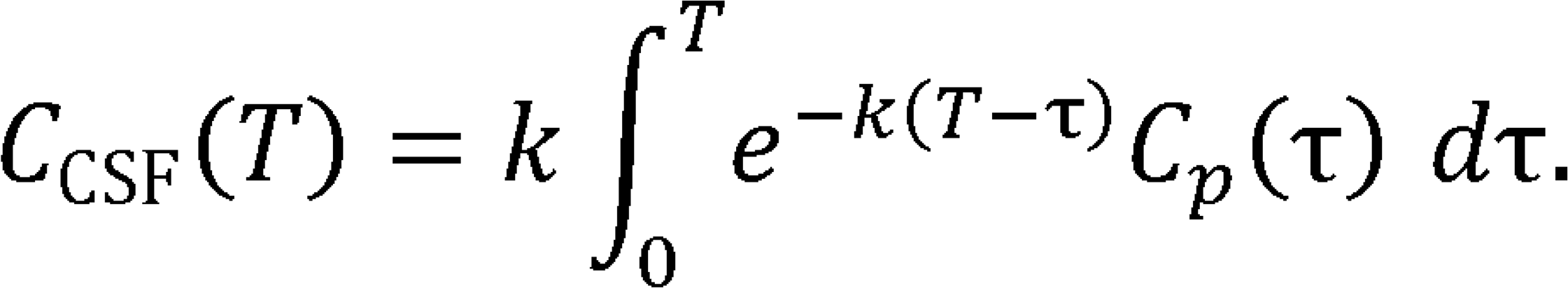

Plasma D2O concentration (C_P_) is measured as the signal changes in the superior sagittal sinus during and after intravenous D2O infusion. Plasma concentration curve was approximated as continuous piecewise-linear function using experimentally measured slopes across infusion epochs. Each segment was integrated analytically to yield the weighted plasma integral *I(k)*. The turnover constant (*k*) was then estimated by numerically solving the nonlinear equation *C_CSF_*(*T*) = *kl*(*k*) using Brent’s method, as implemented in the *uniroot* function in R (v4.3.1).

### Statistical analysis

All analyses were conducted in R (v4.3.1). Global CSF signal changes and cortical signal changes from the indirect dynamic deuterium displacement imaging analysis were compared between groups using the Wilcoxon rank-sum test. For the follow-up ROI analysis, the percent change from baseline was evaluated across conditions using the Jonckheere–Terpstra test with the hypothesized order (D2O > D2O/ACZ > H2O; one-tailed). A threshold of p < 0.05 was considered statistically significant.

## Acknowledgement

The authors thank Professor Joseph J. H. Ackerman for advice and helpful discussions. This work was supported in part by the Radiological Society of North America (RSNA) Research Scholar Grant (#RSCH25-253). H.H. was supported by the TOP-TIER training grant (T32-EB021955) during the study period. MRI studies were performed on a 9.4 T small-animal scanner supported by the National Institutes of Health (NIH) Shared Instrumentation Grant (S10OD026913).

## References

1. Rasmussen MK, Mestre H, Nedergaard M. Fluid transport in the brain. Physiol Rev 2022;102:1025–151.

2. Lun MP, Monuki ES, Lehtinen MK. Development and functions of the choroid plexus–cerebrospinal fluid system. Nature Reviews Neuroscience 2015 16:8 2015;16:445–57.

3. MacAulay N, Keep RF, Zeuthen T. Cerebrospinal fluid production by the choroid plexus: a century of barrier research revisited. Fluids and Barriers of the CNS 2022 19:1 2022;19:1–18.

4. Xie L, Kang H, Xu Q, et al. Sleep drives metabolite clearance from the adult brain. Science (1979) 2013;342:373–7.

5. Orešković D, Radoš M, Klarica M. Role of choroid plexus in cerebrospinal fluid hydrodynamics. Neuroscience 2017;354:69–87.

6. Damkier HH, Brown PD, Praetorius J. Cerebrospinal fluid secretion by the choroid plexus. Physiol Rev 2013;93:1847–92.

7. Orešković D, Klarica M. The formation of cerebrospinal fluid: Nearly a hundred years of interpretations and misinterpretations. Brain Res Rev 2010;64:241–62.

8. Karimy JK, Zhang J, Kurland DB, et al. Inflammation-dependent cerebrospinal fluid hypersecretion by the choroid plexus epithelium in posthemorrhagic hydrocephalus. Nature Medicine 2017 23:8 2017;23:997–1003.

9. Lolansen SD, Rostgaard N, Barbuskaite D, et al. Posthemorrhagic hydrocephalus associates with elevated inflammation and CSF hypersecretion via activation of choroidal transporters. Fluids Barriers CNS 2022;19:1–19.

10. Wang Q, Xia X, Zhang H, et al. Targeting modulation of the choroid plexus blood-CSF barrier and CSF hypersecretion via lipid nanoparticle-mediated co-delivery of siRNA and resveratrol. Nature Communications 2025 16:1 2025;16:1–15.

11. Robert SM, Reeves BC, Alper SL, et al. The choroid plexus links innate immunity to CSF dysregulation in hydrocephalus. Cell 2023;186:764–785.e21.

12. Chiu C, Miller MC, Caralopoulos IN, et al. Temporal course of cerebrospinal fluid dynamics and amyloid accumulation in the aging rat brain from three to thirty months. Fluids Barriers CNS 2012;9:1–8.

13. Liu G, Mestre H, Sweeney AM, et al. Direct Measurement of Cerebrospinal Fluid Production in Mice. Cell Rep 2020;33:108524.

14. Delvenne A, Vandendriessche C, Gobom J, et al. Involvement of the choroid plexus in Alzheimer’s disease pathophysiology: findings from mouse and human proteomic studies. Fluids Barriers CNS 2024;21:1–17.

15. Nedergaard M, Goldman SA. Glymphatic failure as a final common pathway to dementia. Science (1979) 2020;370.

16. Zhou C, Zhou Y, Liu L, et al. Progress and recognition of idiopathic intracranial hypertension: A narrative review. CNS Neurosci Ther 2024;30:e14895.

17. Colman BD, Boonstra F, Nguyen MNL, et al. Understanding the pathophysiology of idiopathic intracranial hypertension (IIH): a review of recent developments. J Neurol Neurosurg Psychiatry 2024;95:375–83.

18. Liu G, Ladrón-de-Guevara A, Izhiman Y, et al. Measurements of cerebrospinal fluid production: a review of the limitations and advantages of current methodologies. Fluids and Barriers of the CNS 2022 19:1 2022;19:1–24.

19. Vela AR, Carey ME, Thompson BM. Further data on the acute effect of intravenous steroids on canine CSF secretion and absorption. J Neurosurg 1979;50:477–82.

20. Pollay M, Curl F. Secretion of cerebrospinal fluid by the ventricular ependyma of the rabbit. 101152/ajplegacy196721341031 1967;213:1031–8.

21. Perera C, Cruz R, Shemesh N, et al. Non-invasive MRI of blood-cerebrospinal fluid-barrier function in a mouse model of Alzheimer’s disease: a potential biomarker of early pathology. Fluids Barriers CNS 2024;21:1–15.

22. Evans PG, Sokolska M, Alves A, et al. Non-Invasive MRI of Blood–Cerebrospinal Fluid Barrier Function. Nature Communications 2020 11:1 2020;11:1–11.

23. Lee H, Ozturk B, Stringer MS, et al. Choroid plexus tissue perfusion and blood to CSF barrier function in rats measured with continuous arterial spin labeling. Neuroimage 2022;261:119512.

24. Bering EA. Water Exchange of Central Nervous System and Cerebrospinal Fluid. J Neurosurg 1952;9:275–87.

25. Mamonov AB, Coalson RD, Zeidel ML, et al. Water and Deuterium Oxide Permeability through Aquaporin 1: MD Predictions and Experimental Verification. Journal of General Physiology 2007;130:111–6.

26. Ibata K, Takimoto S, Morisaku T, et al. Analysis of aquaporin-mediated diffusional water permeability by coherent anti-Stokes Raman scattering microscopy. Biophys J 2011;101:2277–83.

27. Wang FN, Peng SL, Lu CT, et al. Water signal attenuation by D2O infusion as a novel contrast mechanism for 1H perfusion MRI. NMR Biomed 2013;26:692–8.

28. Chen L, Liu J, Chu C, et al. Deuterium oxide as a contrast medium for real-time MRI-guided endovascular neurointervention. Theranostics 2021;11:6240–50.

29. Ackerman JJH, Ewy CS, Becker NN, et al. Deuterium nuclear magnetic resonance measurements of blood flow and tissue perfusion employing 2H2O as a freely diffusible tracer. Proceedings of the National Academy of Sciences 1987;84:4099–102.

30. De Feyter HM, Behar KL, Corbin ZA, et al. Deuterium metabolic imaging (DMI) for MRI-based 3D mapping of metabolism in vivo. Sci Adv 2018;4:7314–36.

31. De Feyter HM, de Graaf RA. Deuterium metabolic imaging – Back to the future. Journal of Magnetic Resonance 2021;326:106932.

32. Urushihata T, Takuwa H, Takahashi M, et al. Distribution of intraperitoneally administered deuterium-labeled water in aquaporin-4-knockout mouse brain after middle cerebral artery occlusion. Front Neurosci 2023;16:1071272.

33. Su S, Qiu Z, Luo C, et al. Accelerated 3D bSSFP Using a Modified Wave-CAIPI Technique with Truncated Wave Gradients. IEEE Trans Med Imaging 2021;40:48– 58.

34. Barbuskaite D, Oernbo EK, Wardman JH, et al. Acetazolamide modulates intracranial pressure directly by its action on the cerebrospinal fluid secretion apparatus. Fluids Barriers CNS 2022;19:1–19.

35. Faraci FM, Mayhan WG, Heistad DD. Vascular effects of acetazolamide on the choroid plexus. Journal of Pharmacology and Experimental Therapeutics 1990;254:23–7.

36. Takamata A, Seo Y, Ogino T, et al. Effects of pCO2 on the CSF Turnover Rate in T1-Weighted Magnetic Resonance Imaging. Jpn J Physiol 2001;51:555–62.

37. Karimy JK, Kahle KT, Kurland DB, et al. A novel method to study cerebrospinal fluid dynamics in rats. J Neurosci Methods 2015;241:78–84.

38. Bito Y, Harada K, Ochi H, et al. Low b-value diffusion tensor imaging for measuring pseudorandom flow of cerebrospinal fluid. Magn Reson Med 2021;86:1369–82.

39. Nazeri A, Hosseini H, Dehkharghanian T, et al. Characterizing the spatial patterns and determinants of cerebrospinal fluid pseudorandom flow in the human brain with low b-value diffusion MRI. Imaging Neuroscience 2025;3:2025.

40. Bulat M, Lupret V, Orehković D, et al. Transventricular and transpial absorption of cerebrospinal fluid into cerebral microvessels. Coll Antropol 2008;32:43–50.

41. Nicholls MG. Independence of the central nervous and the peripheral renin-angiotensin systems in the dog. Hypertension 1979;1:228–34.

42. Selby JB, Mathis JE, Berry CF, et al. Effects of isotonic saline solution resuscitation on blood coagulation in uncontrolled hemorrhage. Surgery 1996;119:528–33.

43. Mueller PJ, Hasser EM. Enhanced sympathoinhibitory response to volume expansion in conscious hindlimb-unloaded rats. J Appl Physiol (1985) 2003;94:1806–12.

44. Barbuskaite D, Oernbo EK, Wardman JH, et al. Acetazolamide modulates intracranial pressure directly by its action on the cerebrospinal fluid secretion apparatus. Fluids Barriers CNS 2022;19:1–19.

45. Bangerter NK, Hargreaves BA, Vasanawala SS, et al. Analysis of multiple-acquisition SSFP. Magn Reson Med 2004;51:1038–47.

46. Avants BB, Tustison NJ, Song G, et al. A reproducible evaluation of ANTs similarity metric performance in brain image registration. Neuroimage 2011;54:2033–44.

47. Avants BB, Tustison NJ, Wu J, et al. An open source multivariate framework for N-tissue segmentation with evaluation on public data. Neuroinformatics 2011;9:381– 400.

48. Jenkinson M, Beckmann CF, Behrens TEJ, et al. FSL. Neuroimage 2012;62:782– 90.

49. Yushkevich PA, Gao Y, Gerig G. ITK-SNAP: An interactive tool for semi-automatic segmentation of multi-modality biomedical images. Proceedings of the Annual International Conference of the IEEE Engineering in Medicine and Biology Society, EMBS 2016;2016-October:3342–5.

50. Perera C, Cruz R, Shemesh N, et al. Non-invasive MRI of blood-cerebrospinal fluid-barrier function in a mouse model of Alzheimer’s disease: a potential biomarker of early pathology. Fluids Barriers CNS 2024;21:1–15.

51. Daoust A, Dodd S, Nair G, et al. Transverse relaxation of cerebrospinal fluid depends on glucose concentration. Magn Reson Imaging 2017;44:72–81.

52. Dobre MC, Marjanska M, Ugurbil K. Blood T1 measurement at high magnetic field strength. Proceedings of the 13th annual ISMRM, Miami 2005. [Epub ahead of print].

